# BridgeBP: A Toolbox for Bridging Brain Parcellations and Standardizing Structural Connectivity Matrices

**DOI:** 10.64898/2026.04.17.718823

**Authors:** Ziqian Zhang, Alex Haopeng Liu, Zhengwu Zhang

## Abstract

Brain network analysis has emerged as a critical framework for understanding the complex organization and function of the human brain, underpinning insights into cognition, behavior, and neuropsychiatric conditions. Central to this approach is the parcellation of the brain into discrete regions, which simplifies high-dimensional connectome data and facilitates the investigation of network architectures. However, the proliferation of brain parcellation schemes introduces significant challenges: different parcellations often yield varying network sizes and measures, complicating cross-study comparisons and the reproducibility of findings. Moreover, most connectome construction pipelines are rigid, typically outputting connectivity matrices from only one or a few parcellation schemes, which limits flexibility. In this paper, we address these issues by introducing BridgeBP, a novel toolbox designed to bridge brain parcellations by leveraging continuous brain connectivity concepts. BridgeBP transforms structural connectivity matrices derived from one parcellation scheme into matrices corresponding to more than 40 alternative schemes, standardizing analyses and enhancing the robustness of network studies. Through extensive evaluations, we demonstrate that BridgeBP enables consistent network comparisons across diverse parcellation frameworks, paving the way for more reproducible and generalizable insights in brain connectome research.

## Introduction

The human brain operates through intricate patterns of communication across distributed neural regions, forming complex networks that underpin cognition, behavior, and emotion (1; 2; 3). Mapping and analyzing these brain networks, collectively known as the connectome, using non-invasive neuroimaging techniques like MRI has become a central focus in modern neuroscience (4; 5; 6). A critical step in connectome analysis involves parcellating the brain, particularly the cerebral cortex, into a set of discrete regions or parcels (7). This parcellation process simplifies the high-dimensional, spatially complex connectome data into a manageable network representation—typically a connectivity matrix where nodes represent brain regions and edges represent the strength of structural or functional connections between them.

Despite great progress, however, the pursuit of understanding brain organization through parcellation faces a significant challenge: the lack of a single, universally adopted parcellation scheme (8; 9; 10; 11). Numerous atlases exist, varying widely in their methodologies (derived using either anatomical landmarks, functional homogeneity, or structural connectivity patterns), scale (number of parcels), and spatial boundaries (12; 13). This proliferation, while reflecting diverse research goals and perspectives, introduces substantial variability into connectome analysis. Studies have shown that the choice of parcellation can significantly alter the resulting network structure, graph theoretical measures, and downstream statistical inferences, thereby complicating cross-study comparisons and hindering the reproducibility of findings (14; 15). Furthermore, most existing connectome construction pipelines are designed with limited flexibility, often hardcoded to generate outputs based on only one or a small selection of pre-defined atlases (16; 17). Adapting these pipelines to accommodate different parcellation schemes is often non-trivial, requiring significant reprocessing of raw data, specialized expertise, and introducing potential for errors. While valuable initiatives exist to aggregate and standardize brain atlases themselves (18; 19), these resources do not directly address the critical need to transform connectivity results derived from one parcellation into another. This gap limits the ability to integrate data and findings across studies.

To address the limitations of parcellation-dependent connectome analyses, we present BridgeBP (Bridging Brain Parcellations), a computational toolbox that standardizes structural connectivity matrices across diverse brain atlases. BridgeBP tackles the parcellation variability problem by transforming a connectivity matrix derived from one atlas into an equivalent representation under another atlas. Drawing on the concept of continuous connectivity representations, which model connectivity across the entire cortical surface rather than discrete region pairs (20; 21; 22), BridgeBP first estimates a high-resolution continuous connectivity profile from the source atlas-based matrix. It then integrates this profile over the regions defined by the target atlas to construct the corresponding discrete connectivity matrix. This framework enables seamless conversion between over 40 widely used brain parcellations included in the toolbox. Figure 1 illustrates the BridgeBP pipeline’s workflow.

**Figure 1.**
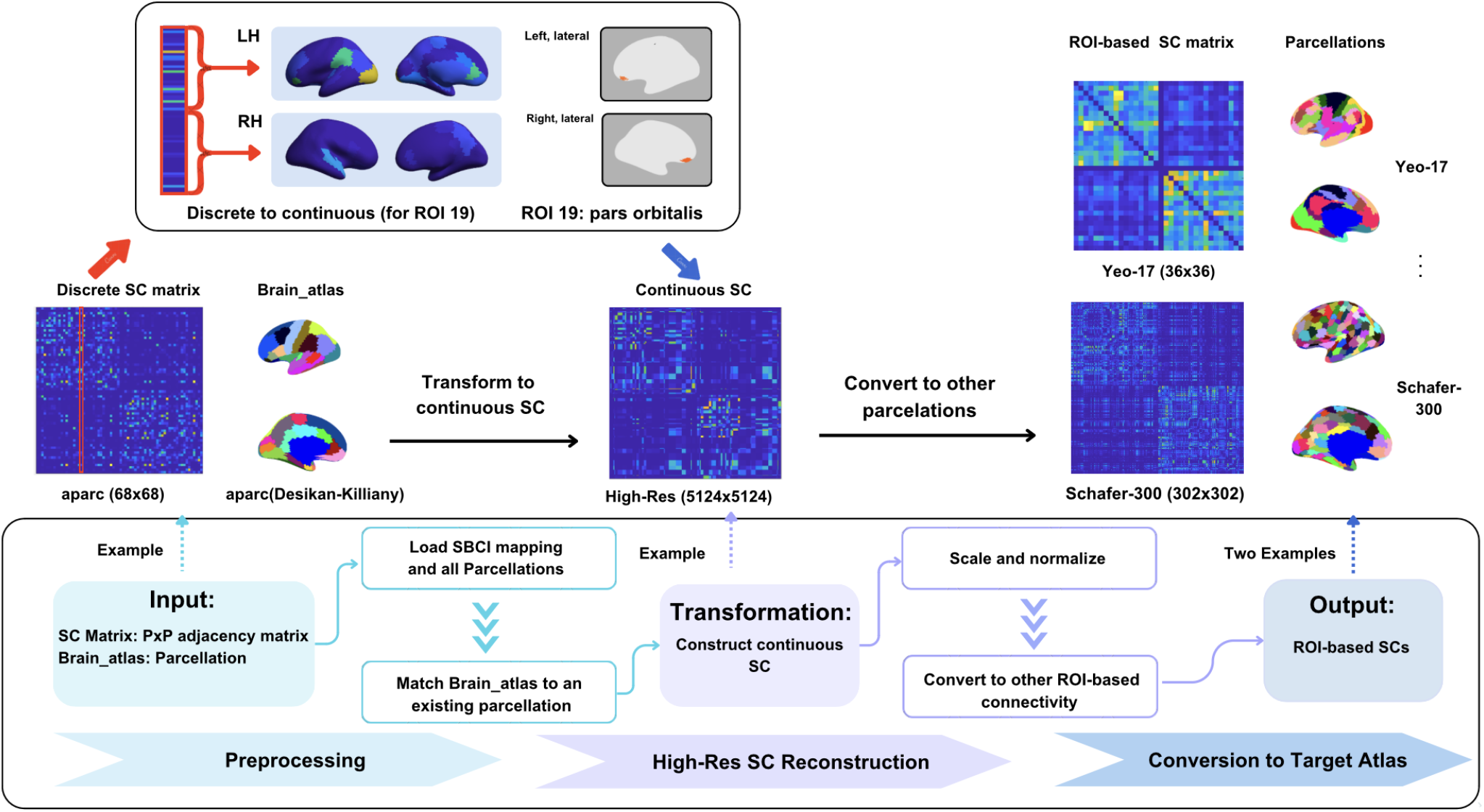
Workflow of the BridgeBP toolbox. The pipeline takes a discrete structural connectivity (SC) matrix defined on one brain atlas as input and transforms it into discrete SC matrices aligned with any target atlas. By leveraging high-resolution surface mapping and interpolation, the toolbox enables flexible SC transformation across parcellations of varying resolution and type. Note that the examples shown involve the Desikan–Killiany atlas (23), the Yeo atlas Yeo-17 (13), and the Schaefer atlas Schafer-300 (10). ROI stands for region of interest, and SBCI refers to surface-based connectome integration.

By providing the atlas conversion function, BridgeBP offers several key advantages for current brain network studies: (1) It facilitates direct comparisons of network studies using different parcellations. (2) It enhances the reproducibility and generalizability of connectome research by allowing results to be expressed in multiple atlas frameworks. (3) It increases analytical flexibility, enabling researchers to explore how network properties manifest across different parcellation choices without extensive reprocessing. (4) It includes tools for comparing and visualizing matrices across parcellation schemes, further aiding interpretation. The BridgeBP toolbox is available in https://github.com/sbci-brain/BridgeBP.git with detailed documentation.

## Results

### Atlases Available for Conversion

BridgeBP incorporates an extensive collection of 44 diverse brain atlases, organized to cover multiple resolutions and methodologies of cortical parcellation. These atlases are grouped broadly based on their derivation methods: anatomical, functional, connectivity-based, and multi-modal. Anatomical atlases include widely-used ones such as the Desikan–Killiany atlas (23) with 68 cortical regions, the Destrieux atlas (24) containing 148 cortical regions, and variants from the PALS-B12 family (25) with resolutions ranging from 10 to 82 regions. Functional atlases primarily derived from resting-state fMRI include the Gordon atlas (9) delineating 333 cortical regions, the Yeo atlas (13) available in 7- and 17-network versions, and the Schaefer atlas series (10) covering multiple granularities from 100 to 1000 regions. Multi-modal atlases, integrating anatomical, functional, and other MRI modalities, include the Brainnetome atlas (26) with 210 cortical regions and the HCP-MMP1 atlas (12) consisting of 360 cortical areas. Additionally, BridgeBP includes the structural connectivity-based CoCoNest atlas family (11), which uniquely leverages hierarchical structural connectivity information. This atlas family is generated using an agglomerative clustering approach that constructs a complete binary hierarchy, followed by a pruning procedure optimizing the balance between complexity and fit. The resulting parcellations range from 9 to 500 regions, providing a detailed, multiscale exploration of structural connectivity patterns.

Table 1 details the 44 atlases in BridgeBP. Collectively, these atlases offer comprehensive coverage of parcellation strategies employed in network neuroscience. By integrating diverse methodological approaches and multiple resolution levels, BridgeBP facilitates robust, comparative, and standardized assessments of brain connectivity across research studies.

**Table 1.** This table contains the atlases included in BridgeBP, along with the number of regions of interest (ROIs), a brief description of each atlas, and the classification of atlas type as anatomical-, functional-, connectivity-, or multi-modal-based.

| Atlas Family | # ROIs (per version) | # versions | Description | Type |
| --- | --- | --- | --- | --- |
| Brainnetome | 210 | 1 | Fine-grained parcellation with 210 cortical regions (26). | Multi-modal-based |
| Gordon | 333 | 1 | Parcellation derived from resting-state functional connectivity (RSFC) boundary mapping. Identifies putative cortical areas by detecting abrupt RSFC transitions. Produces parcels with high functional homogeneity and reliable network structure (9). | Functional-based |
| HCP | 360 | 1 | HCP MMP 1.0 has been built using surface-based registrations of multimodal MR data from 210 HCP subjects. 180 areas per hemisphere have been identified (12). | Multi-modal-based |
| Cortical Hubs | 14 | 1 | Defined by cortical thickness differences between older controls and Alzheimer’s patients using OASIS data. Developed to highlight regions vulnerable to AD-related atrophy. Provided as regions of interest in supplemental materials (27). | Anatomical-based |
| Yeo | [7; 17] | 2 | Parcellations from 1000 subjects. Clustering revealed 7 and 17 functional networks across cortex, including sensory-motor and association circuits (13). | Functional-based |
| Desikan-Killiany & Destrieux | [68; 148; 155] | 3 | Automatic cortical labeling using surface geometry and training data. Includes Desikan-Killiany (23; 28) and Destrieux (“aparc.a2005s”, “aparc.a2009s”) (24). Parcellations define gyral and sulcal regions based on anatomical conventions and curvature-derived boundaries. | Anatomical-based |
| PALS-B12 | [10; 18; 48; 82] | 4 | A population-average surface-based atlas from 12 adults, built with landmark-constrained registration. It supports sulcal alignment, multi-fiducial mapping, and reveals cortical asymmetries. Includes Brodmann, Lobes, Orbitofrontal, and Visuo-topic ROIs on fsaverage (25). | Anatomical-based |
| Schaefer | [100-1000] | 10 | Parcellations based on resting-state fMRI from 1489 subjects using surface-based alignment and a gradient-weighted Markov random field. Extends Yeo networks (13) by locally refining functional boundaries (10). | Functional-based |
| CoCoNest | [9-500] | 21 | A multiresolution family of structural parcellations derived from high-resolution connectivity data. Generated using agglomerative clustering and pruning. Enables exploration of the nested organization of the structural connectome across multiple spatial scales (11). | Connectivity-based |

### Cross-Atlas Conversion Consistency

Our toolbox, BridgeBP, is designed to transform SC matrices between different brain atlas representations. This function is achieved through an intermediate step: the generation of a high-resolution, continuous representation of SC (referred to as continuous SC, visualized in Figure 1 from an initial discrete, atlas-based SC). To rigorously assess the fidelity of these transformations, we first performed two key evaluations using data from 100 subjects from the Adolescent Brain Cognitive Development Study (ABCD) (5). For comparison, “ground truth” continuous SC and corresponding discrete atlas-based SCs were derived using the SBCI pipeline (20). Here, “ground truth” SC refers to the continuous SC directly obtained from the SBCI pipeline and used as the reference for evaluating reconstruction fidelity. Importantly, BridgeBP is compatible with SC matrices obtained from parcellation schemes in Table 1 and is not restricted to SBCI-derived SCs.

First, we evaluated the accuracy of reconstructing the continuous SC representation. Starting from various discrete atlas-based SCs (each with a different number of parcels), we used our approach to generate a corresponding continuous SC. This reconstructed continuous SC was then compared against the ground truth continuous SC. Figure 2 presents these comparisons, using Pearson correlation computed over non-zero SC entries as the metric of similarity. Supplementary Figure S3 shows the scatter plots of true versus reconstructed SC for selected parcellations. The results show that the fidelity of the reconstructed continuous SC improves as the number of parcels in the source atlas increases. This indicates that richer, more detailed source parcellations enable a more accurate estimation of the underlying continuous connectivity landscape.

**Figure 2.**
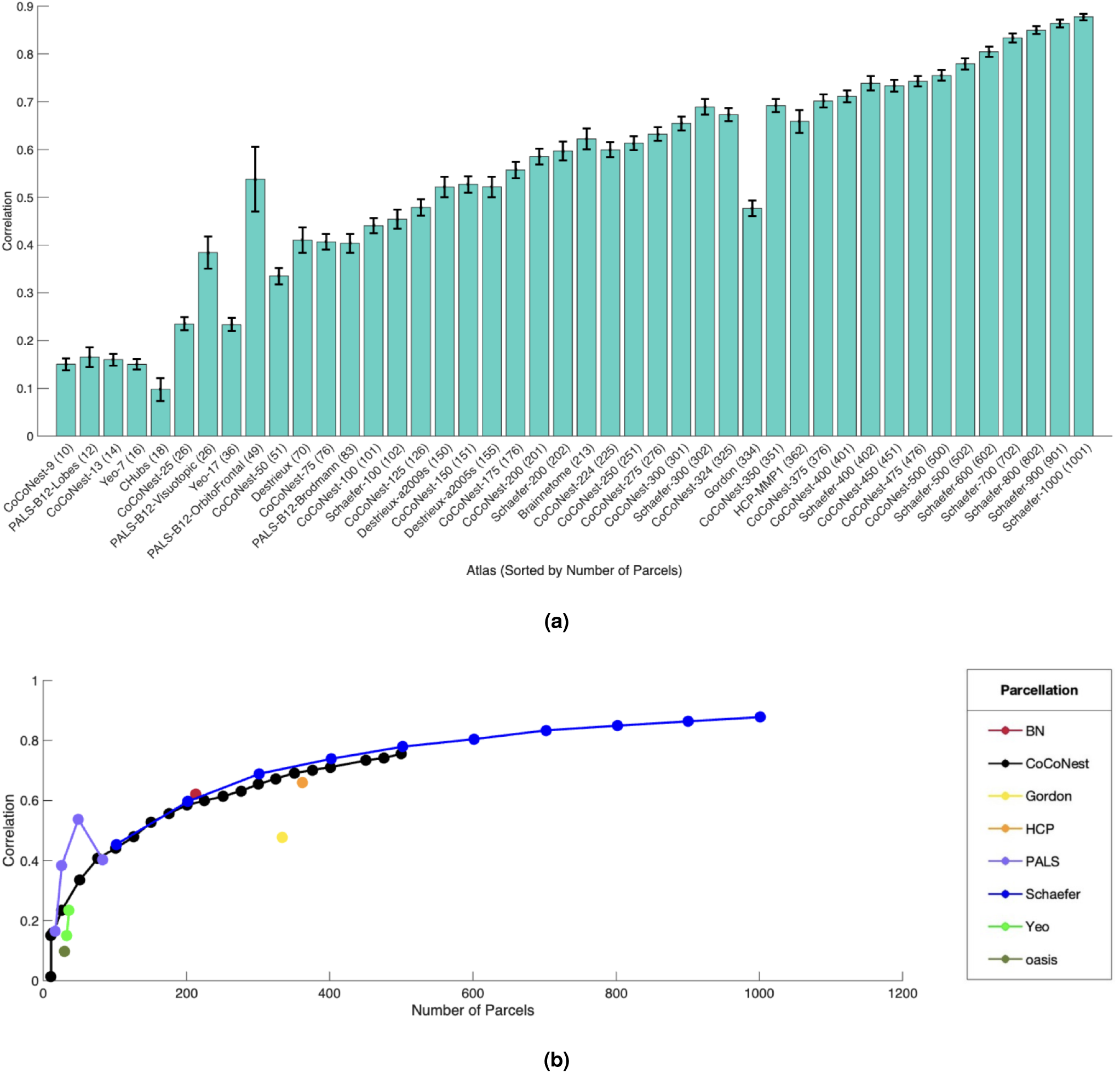
Correlations between the reconstructed and ground truth continuous SC over non-zero entries. (A) Mean correlations with standard deviations across 100 subjects for each brain atlas. Correlation quantifies how well the reconstructed continuous SC matches the ground truth SC. (B) Relationship between the number of parcels and SC reconstruction performance across atlas classes.

Second, we assessed the accuracy of converting SC data from one discrete atlas directly to another (termed ‘cross-atlas conversion’). For a given source atlas-based SC, BridgeBP converts it into the format of a target atlas. The accuracy of this cross-atlas conversion was then evaluated by comparing the newly generated target atlas SC against its “ground truth” counterpart (i.e., the SC for the target atlas derived directly from the ground truth continuous SC). Figure 3 displays the mean correlation results. A clear pattern emerged: converting from a low-resolution discrete SC (fewer parcels) to a high-resolution discrete SC (more parcels) resulted in diminished accuracy. Conversely, converting from a high-resolution discrete SC to a low-resolution one yielded nearly perfect reconstructions. This asymmetry is intuitive: a higher-resolution initial SC allows for a more precise estimation of the intermediate continuous SC, which in turn facilitates more accurate subsequent conversions to other atlas-based SCs, especially when the target parcellation is coarser. Additional metrics for this experiment, including accuracy measured by Mean Squared Error (MSE) and the standard deviation of the correlations, are presented in Supplementary Figure S2.

**Figure 3.**
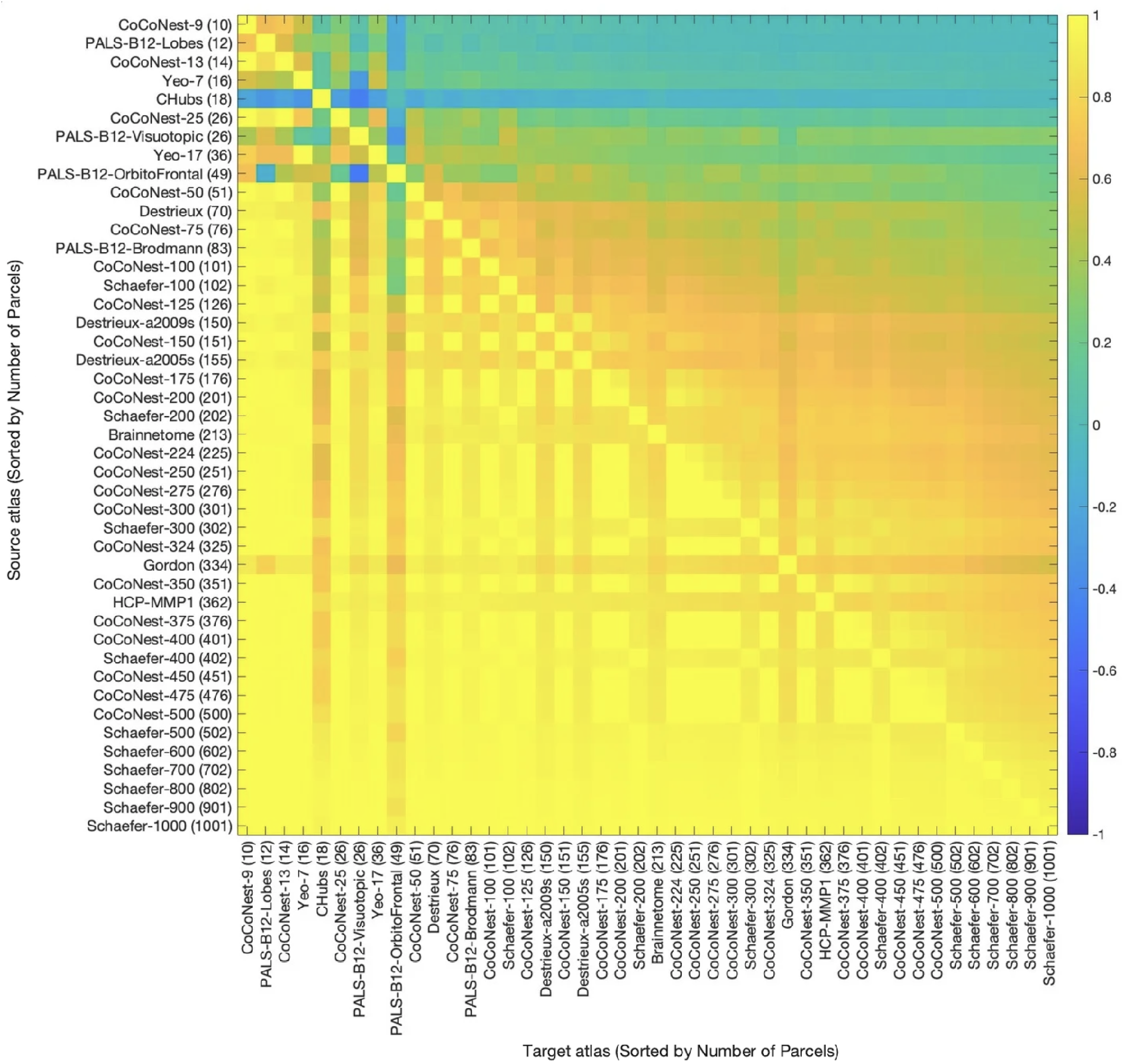
Mean conversion accuracy (measured using Pearson correlation) matrices. For each source atlas-based SC, BridgeBP was used to convert it to various atlas-based SCs. The conversion accuracy was measured between the converted and ground truth SCs. Row names indicate the target atlas, and column names indicate the source atlas.

To complement the SBCI-based evaluation, we also tested BridgeBP on an independent dataset of 100 subjects from the Human Connectome Project (HCP), which provides Desikan-68 and Destrieux-148 SC matrices generated using a different SC estimation pipeline, named PSC pipeline(16). Because these SCs were produced outside the SBCI framework and do not include continuous SC ground truth, we evaluated the pairwise atlas conversions supported by the available data. Specifically, because the PSC pipeline provides empirical SC matrices for both atlases for every subject, we utilized these paired matrices to establish source inputs and target ground truths. We applied BridgeBP to transform a subject’s PSC-generated SC matrix from a source atlas (e.g., Desikan-68) into a target atlas (e.g., Destrieux-148), and subsequently evaluated the accuracy by comparing this BridgeBP-predicted matrix directly against the subject’s actual PSC-generated matrix in the target atlas. This additional experiment demonstrates that BridgeBP can operate on SCs derived from other preprocessing pipelines and is not restricted to SBCI-generated data. Table 2 summarizes the HCP results, showing the mean correlation and normalized squared error (NSE) for conversions between the Desikan and Destrieux atlases. BridgeBP achieves good conversion accuracy on these independently generated SCs, with higher performance when converting from the finer Destrieux atlas to the coarser Desikan atlas.

**Table 2.** Mean correlation and normalized squared error (NSE) of atlas conversion performance on HCP SC data. Results are averaged across 100 subjects, comparing transformations between Desikan and Destrieux parcellations.

| Conversion | Mean $r$ | NSE |
| --- | --- | --- |
| Desikan → Destrieux | 0.7516 | 1.49 |
| Destrieux → Desikan | 0.8374 | 0.645 |

### Application to Pre-trained Machine Learning Models

We demonstrate one important application of BridgeBP in standardizing connectome matrices. A significant challenge in leveraging pre-trained machine learning models for connectome analysis is the common requirement for input data to conform to a specific brain atlas format. If a researcher’s SC data are parcellated using a different atlas than the one on which a model was trained, direct application of that pre-trained model is often impossible. BridgeBP addresses this critical gap by enabling the transformation of SC matrices between different parcellations.

To demonstrate this application, we evaluated the utility of BridgeBP in an SC-based classification task. We employed four representative neural network architectures (29): Graph Convolutional Network (GCN), Graph Isomorphism Network (GIN), Subgraph-based GNN (Moment-GNN), and a Multilayer Perceptron (MLP). The objective was to classify gender based on their SC matrices. For this experiment, the HCP360 atlas was considered the “target” atlas, meaning that the machine learning (ML) models were intended to work with SC data in this format (representing the “ground truth” parcellation for the model). We then simulated scenarios where SC data were initially available in different atlases: For the ABCD data, Schaefer-1000, Schaefer-500, and Schaefer-100, representing high-, mid-, and low-resolution parcellations relative to HCP360. For the HCP dataset, we additionally included Desikan-68 and Destrieux-148 SC matrices derived using other SC extraction pipeline(16), representing SC data generated outside the SBCI framework. BridgeBP was used to convert all of these source SC matrices into the HCP360 format.

We compared model performance under several conditions (See Table 3 and 4): (a). Ground Truth: Models trained and tested directly on SC matrices derived using the HCP360 atlas; (b). Pre-trained with BridgeBP conversion: Models trained on HCP360 SC matrices, then tested on BridgeBP converted HCP360 SC data; And (c). Fine-tuned with BridgeBP converted data: Models from Condition (a) were subsequently fine-tuned using a portion of the BridgeBP converted HCP360 SC data before being tested on a separate BridgeBP converted test set.

**Table 3.**
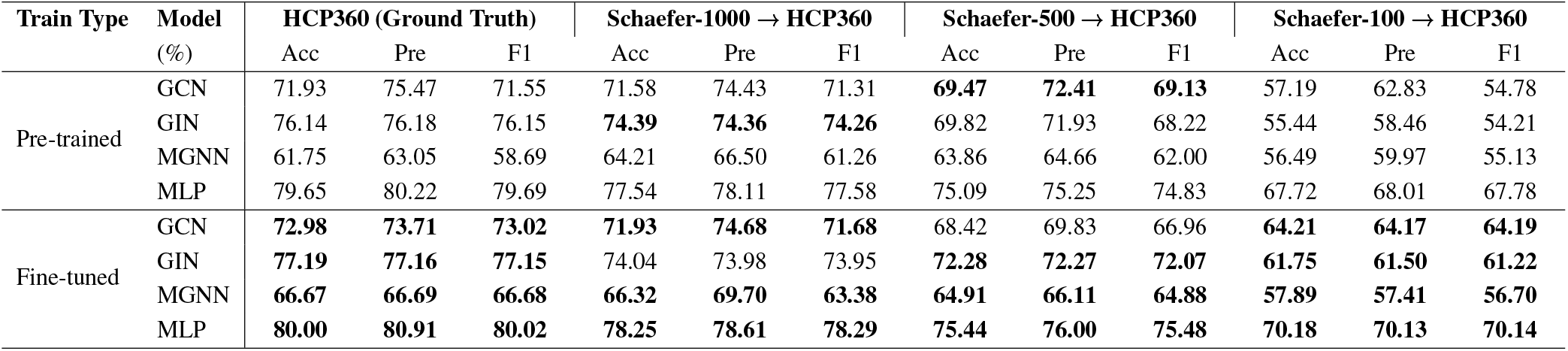
Classification performance (Accuracy, Precision, F1-score, in %) of machine learning models on ABCD SC data. Models trained on ground truth HCP360 data were applied on data converted by BridgeBP from Schaefer atlases (Schaefer-1000, -500, -100) to HCP360, with and without subsequent fine-tuning on the converted data.

**Table 4.** Similar to Table 3, but with HCP data. The SC matrices were generated by PSC pipeline, serving as an independent validation of BridgeBP’s ability to transfer atlases for other SC extraction pipelines. Models trained on ground truth HCP360 data under the ABCD dataset were evaluated on the converted datasets, with and without fine-tuning on the converted SC matrices.

| Train Type | Model (%) | Desikan-68 → HCP360 | | | Destrieux-148 → HCP360 | | |
| --- | --- | --- | --- | --- | --- | --- | --- |
|  |  | Acc | Pre | F1 | Acc | Pre | F1 |
| Pre-trained | GCN | 52.39 | 48.01 | 38.05 | 49.56 | 49.59 | 49.57 |
|  | GIN | 52.74 | 51.48 | 38.51 | 51.50 | 49.96 | 46.66 |
|  | MGNN | 52.74 | 27.82 | 36.43 | 52.74 | 27.82 | 36.43 |
|  | MLP | 52.74 | 27.82 | 36.43 | 52.74 | 27.82 | 36.43 |
| Fine-tuned | GCN | **66.55** | **66.75** | **66.57** | **66.02** | **66.92** | **65.90** |
|  | GIN | **66.90** | **66.90** | **66.90** | **69.03** | **69.73** | **68.98** |
|  | MGNN | **61.42** | **61.93** | **60.09** | **63.19** | **63.11** | **63.08** |
|  | MLP | **70.80** | **70.91** | **70.81** | **70.80** | **71.32** | **70.78** |

The results (Table 3 and 4) show that SC matrices transformed by BridgeBP are compatible with various deep learning models and support the pre-training and fine-tuning pipeline for brain network analysis. For instance, when using SC data initially from the Schaefer-1000 atlas and converted to HCP360 by BridgeBP, the MLP model achieved an accuracy of 77.54% on the pre-trained model. It was improved to 78.25% after fine-tuning. This is reasonably close to the MLP performance on ground truth HCP360 data (i.e., 79.65%). We see that fine-tuning consistently improved model accuracy across datasets and atlases.

## Discussion

This study introduced BridgeBP, a computational toolbox for transforming SC matrices across over 40 diverse brain parcellation schemes. By enabling interoperability between studies using different atlases, BridgeBP addresses a key challenge in network neuroscience. We demonstrated its conversion consistency and its practical utility in facilitating ML applications with SC data.

### Key Achievements and Implications

BridgeBP enables the conversion of SC matrices between different parcellations by employing an intermediate high-resolution continuous SC representation. Notably, the tool operates directly on atlas-based SC matrices, allowing users to bypass reprocessing of raw neuroimaging data when such matrices are already available. Our evaluations (Figures 2 and 3) validated this approach, showing that conversion fidelity increases with source atlas resolution and that high-to-low resolution transformations achieved strong performance. This functionality is vital for harmonizing diverse datasets, thereby facilitating more direct and reliable comparisons of neurobiological findings by reducing atlas-specific methodological biases. We have also demonstrated an important practical use of BridgeBP in using and fine-tuning pre-trained ML models for SC data. As shown in Table 3 and 4, SC data converted by BridgeBP to match a model’s required atlas format (e.g., from Schaefer, Desikan, or Destrieux to HCP360) can yield classification performance that is almost comparable to models trained on ground-truth target-atlas data, especially with fine-tuning. This democratizes the use of BridgeBP for studies with disparately parcellated data, promoting data reuse and reproducibility in computational neuroscience.

### Challenges and Limitations

Despite its utility, BridgeBP faces limitations inherent to atlas-level representations of SC. The current BridgeBP assumes that connectivity within each ROI pair is uniformly distributed. As a result, the intermediate continuous SC recovered from a discrete SC becomes piecewise constant (Equation 2). Therefore, fine-scale spatial variations lost during parcel-level aggregation cannot be fully recovered, resulting in reduced fidelity when converting from low-resolution to high-resolution atlases (Figure 3). This limitation reflects a fundamental constraint of working with ROI-based SC rather than a shortcoming of the algorithm itself. BridgeBP is designed as a principled, interpretable framework that avoids introducing dataset-specific artifacts by relying solely on the information present in the source SC. However, improving the recovery of fine-grained structure remains an important direction for future development. Additionally, the current implementation operates on streamline-count connectivity features and does not yet support other measures such as mean fractional anisotropy.

### Future Directions and Opportunities

BridgeBP opens several promising avenues for future network neuroscience research. A significant opportunity lies in leveraging BridgeBP to harmonize large-scale, existing SC datasets, thereby creating a unified dataset for training sophisticated brain network foundation models and generative models capable of learning fundamental connectivity patterns. Concurrently, extending BridgeBP’s capabilities beyond streamline counts to transform other structural connectivity metrics (e.g., mean fractional anisotropy) and adapting it for functional connectivity data will be crucial for comprehensive multi-modal investigations. Our results (Supplementary Figure S4) also indicate that smoothing the intermediate continuous SC 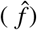 may help improve transformations from coarse to finer parcellations. We employed a Riemannian diffusion kernel (30) to smooth the continuous SC on the cortical surface. Moderate smoothing can enhance reconstruction accuracy for low-resolution atlases, whereas excessive smoothing may oversmooth fine-scale structure in higher-resolution data. Future work will explore adaptive or data-driven smoothing strategies that selectively regularize coarse parcellations while preserving detailed connectivity patterns. The intermediate continuous SC also offers a unique basis for informing connectome densification strategies, potentially yielding richer inputs for machine learning models and improving predictive power for clinical or cognitive outcomes. Finally, expanding BridgeBP to encompass a broader range of brain structures, including subcortical regions and the cerebellum, will be essential for whole-brain analyses and will further solidify its utility as a standard connectivity conversion toolbox.

## Methods

### Brain Connectome Data Compilation

The primary data for this study were sourced from the Adolescent Brain Cognitive Development Study (ABCD) (5), one of the largest publicly available neuroimaging datasets. A total of 2529 subjects were included in our analysis, of whom 1273 (50.34%) were female. Details about imaging acquisition can be found in (5). The diffusion MRI (dMRI) and T1-weighted (T1w) MRI data were processed using the Surface-based Connectivity Integration (SBCI) pipeline (20) to generate high-resolution continuous connectivity matrices. The SBCI pipeline preprocessing steps include standard procedures such as skull stripping, bias field correction, intensity normalization, and cortical surface reconstruction for T1w MRI. For dMRI, SBCI first estimates the fiber orientation distribution function (fODF) at the voxel level and then uses the surface-enhanced tractography (SET) algorithm (31) to extract white matter fiber tracts, forming structural connectivity between different brain regions. SET leverages white surface geometry to enhance fiber tracking accuracy and ensures that the estimated streamline fiber tracts end on the cortical surface.

Let Ω denote the cortical surface and let 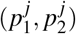 denote a pair of endpoints for a white-matter fiber tract estimated with SET. Since SET ensures 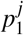 and 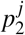 are on the cortical surface, we obtain pairs of endpoints on Ω to represent the structural brain connectivity. Denote this set of observed endpoints as 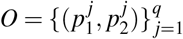, SBCI models *O* as a realization of a spatial point process on the product space Ω× Ω. The underlying structure of this process is characterized by an unknown, non-negative intensity function *u* : Ω× Ω → [0, ∞). This intensity function *u*(*ω*_1_, *ω*_2_) relates to the expected number of streamline endpoints falling within infinitesimal regions around *ω*_1_ and *ω*_2_. Specifically, for any two measurable regions *E*_1_, *E*_2_ ⊂ Ω, the expected number of streamlines connecting *E*_1_ and *E*_2_, denoted #(*E*_1_, *E*_2_), is given by:

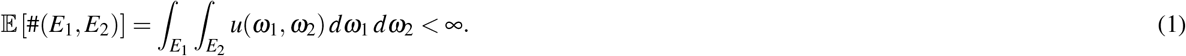

The total expected number of streamlines is *E*[*q*] = _Ω Ω_ *u*(*ω*_1_, *ω*_2_) *dω*_1_ *dω*_2_. The object often targeted for estimation and comparison is the normalized version of *u*, the probability density function *f* (*ω*_1_, *ω*_2_):

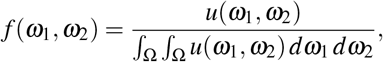

assuming the total intensity is positive and finite. This density *f* represents the continuous structural connectome, capturing the probability distribution of streamline connections across the cortical surface product space. By construction, *f* is symmetric, i.e., *f* (*ω*_1_, *ω*_2_) = *f* (*ω*_2_, *ω*_1_). SBCI saves *f* as a matrix of 5124 × 5124 based on a sparse grid on Ω (20).

### Transformation Process

Let 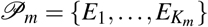 be a cortical parcellation of Ω (e.g., the *m*-th parcellation in BridegeBP), where *K*_*m*_ denotes the total number of parcels, and let 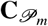 be the SC matrix formed by the parcellation, where 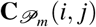 (and 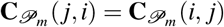) represents the connection strength between the parcel *E*_*i*_ and *E*_*j*_ in 𝒫_*m*_. In this paper, we consider the connection strength measure as the number of streamlines (or fiber tracts) between two parcels. The total number of streamlines is given by 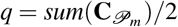. In BridgeBP, we have collected 44 different atlases, so the atlas index *m* goes from 1 to 44. **Our goal is to estimate a continuous connectivity** 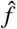 **from the input discrete adjacency matrix** 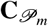 **and the atlas** 𝒫_*m*_. Below, we describe the details of our method.

The surface area of region *E*_*i*_ is given by: 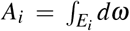. A piecewise-constant function 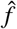 is then constructed on Ω × Ω by assigning

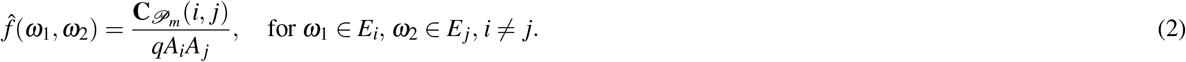

This function is symmetric and constant on each region pair *E*_*i*_ × *E*_*j*_. Since we do not consider self-connection, the diagonal *ω*_1_ = *ω*_2_ is set to 0. The function 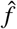 satisfies the integral constraint:

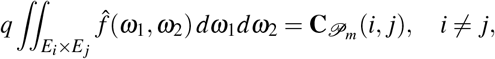

which provides a continuous representation of structural connectivity used in the SBCI pipeline.

Now given a different parcellation 𝒫_*m*′_, we compute the estimated connectivity matrix 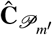 for this new parcellation. Let the new parcellation be denoted as 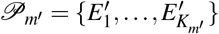, where *K*_*m*′_ is the number of parcels in this atlas. Our goal is to estimate the number of streamlines connecting region 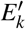 and region 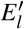, denoted by 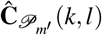, using the piecewise-constant function 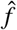 derived from the source parcellation 𝒫_*m*_. Then 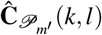 can be computed by integrating 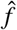 over the product region 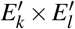 and scaling by the total number of streamlines *q*. That is:

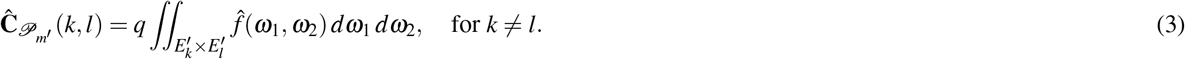

Equation (3) provides the mechanism to transform the streamline counts from one parcellation 𝒫_*m*_ to another 𝒫_*m*′_ using the continuous connectivity as a bridge. The resulting matrix 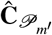 is symmetric due to the symmetry of 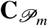 and 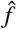.

### Cross-Atlas Conversion Consistency Evaluation

To quantitatively assess the performance and consistency of the SC transformations within BridgeBP, we performed two primary evaluations using *N* = 100 subjects selected from the ABCD dataset. These evaluations leveraged the ground truth continuous SC, *f*_*i*_ (the 5124 × 5124 matrix representation generated by the SBCI pipeline for subject *i*), and ground truth discrete SC matrices, 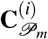, derived by parcellating *f*_*i*_ using each atlas 𝒫_*m*_.

#### 1. Accuracy in Recovering Continuous SC

First, we evaluated the fidelity of reconstructing the continuous SC representation from an initial discrete SC matrix, 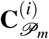, associated with a source atlas 𝒫_*m*_. For each subject *i* and each source atlas 𝒫_*m*_, the estimated continuous SC 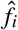 was computed using Equation (2) based on the input 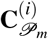. This reconstructed 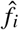 was then directly compared to the subject’s ground truth continuous SC, *f*_*i*_. The accuracy was quantified using the Pearson correlation coefficient between the vectorized high-resolution matrix representations of 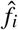 and *f*_*i*_. The results, illustrating how reconstruction accuracy varies with the resolution of the source atlas 𝒫_*m*_ (averaged across *N* subjects), are presented in Figure 2.

#### 2. Accuracy of Cross-Atlas Discrete SC Conversion

Second, we assessed the accuracy of converting an SC matrix from a source atlas 𝒫_*m*_ to a different target atlas 𝒫_*m*′_. For each subject *i*, starting with the ground truth discrete SC matrix 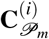 for the source atlas, we first derived the intermediate continuous representation 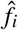 (as per Equation (2)). Then, this 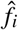 was used to compute the transformed discrete SC matrix for the target atlas, 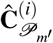, using Equation (3).

The accuracy of this conversion was determined by comparing the transformed matrix 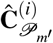 against the ground truth discrete SC matrix for the *target* atlas, 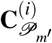 (obtained by parcellating the subject’s ground truth continuous SC, *f*_*i*_, with atlas 𝒫_*m*′_). For each subject *i* and each pair of source (*m*) and target (*m* ′) atlases, the conversion accuracy was quantified by the Pearson correlation coefficient calculated between the upper-triangular elements (excluding the main diagonal) of the respective vectorized matrices.

The mean conversion accuracy across *N* subjects for each atlas pair was then computed, which results in an *M* × *M* matrix (where *M* = 44 is the total number of atlases/versions), where entry (*m, m*′) represents the average accuracy of converting from atlas 𝒫_*m*_ to atlas 𝒫_*m*′_. These mean accuracies are visualized in Figure 3. Furthermore, the standard deviation of the accuracy scores was calculated. This standard deviation matrix (shown in the Supplement) quantifies the inter-subject variability of the conversion accuracy. Low standard deviation alongside high mean correlation indicates a robust and consistent SC conversion process.

### Machine Learning Application

We designed an SC-based classification experiment to evaluate the practical utility of BridgeBP in facilitating the use of pre-trained machine learning models with SC data from disparate atlases.

The SC data for this experiment were derived from a subset of subjects from the ABCD study dataset (*n* = 2,529) (5). The subset was partitioned into training (*n* = 2,149), testing (*n* = 285), and fine-tuning (*n* = 95) cohorts. To ensure that our evaluation of BridgeBP is not limited to SC derived from SBCI, we also included HCP dataset (*n* = 1065). This data was partitioned into testing (*n* = 565), and fine-tuning (*n* = 500) cohorts. Unlike ABCD, the HCP dataset contains SC matrices generated under the Desikan-Killiany (23; 28) and Destrieux (24) parcellations using a different SC estimation pipeline named PSC (16).For each subject in the above subsets, we considered SC matrices generated under different parcellations.

- **Target Atlas (HCP360):** “Ground truth” SC matrices for the classification task were generated by parcellating each subject’s high-resolution continuous SC (obtained via the SBCI pipeline) using the HCP360 atlas (12). These matrices served as the benchmark data for training and testing models in the ideal scenario where the data perfectly match the model’s expected input format. These SC matrices are obtained by parcellating SBCI-derived high-resolution continuous SC into the HCP360 atlas. Note that it does not imply a biological ground truth of SC.
- **Source Atlases (Schaefer Variants):** To simulate scenarios where data are available in a different format, SC matrices were also generated for each subject in the ABCD dataset using three variants of the Schaefer atlas (10): Schaefer-1000, Schaefer-500, and Schaefer-100 parcels.
- **Source Atlases (Desikan and Destrieux)**: To assess generalizability, SC matrices were also generated for a subset of subjects in the HCP dataset using the Desikan (23; 28) and Destrieux (24) parcellations with the PSC pipeline (16). PSC-derived SC matrices were constructed independently from SBCI and were used solely as alternative source-format inputs for BridgeBP conversion.
- **BridgeBP Conversion:** For each subject and each Schaefer source atlas, the corresponding source SC matrix was transformed into the target HCP360 atlas format using BridgeBP. This resulted in three sets of converted HCP360 SC matrices for ABCD and two sets for HCP.

Our classification task was to use SC matrices to predict gender. Four machine learning models were implemented to evaluate the “ground truth” and converted SC data classification performance. These models were chosen to represent complementary architectural families, graph-based networks versus fully connected networks, with differing sensitivities to input structure. Model configurations followed those in Ding et al. (29):

1. **Graph Convolutional Network (GCN)**: A two-layer spectral GCN without residual connections, performing neighborhood aggregation over the SC graph structure.
2. **Graph Isomorphism Network (GIN)**: A two-layer GIN architecture with sum-based aggregation, achieving expressiveness equivalent to the Weisfeiler-Lehman graph isomorphism test.
3. **Moment-GNN (MGNN)**: Topological features (*K* = 10, dimension = 45 each) were extracted from the SC adjacency matrix and concatenated with correlation vectors. A GIN model served as the backbone network.
4. **Multilayer Perceptron (MLP)**: A three-layer fully connected feedforward network that received vectorized SC matrices (flattened lower triangle) as input.

For each of the four ML models and each of the five BridgeBP-converted datasets (three sets of converted HCP360 SC matrices from the ABCD dataset and two from the HCP dataset), as well as the ground truth HCP360 data, we performed the following training and evaluation protocols:

1. **Training on Ground Truth HCP360 Data:** Models were trained using only the SC matrices directly derived from the HCP360 atlas. This serves as the performance benchmark.
2. **Pre-training on BridgeBP-Converted Data:** Models were trained in protocol and evaluated on the test set composed of BridgeBP-converted HCP360 SC matrices.
3. **Fine-tuning after BridgeBP Conversion:** Models that were pre-trained as described in step 1 were subsequently fine-tuned. This involved continuing the training process for a limited number of epochs using a portion of the training data comprised of BridgeBP-converted HCP360 SC matrices. Evaluation was then performed on the test set of BridgeBP-converted HCP360 SC matrices.

Using the training data, we employed 10-fold cross-validation to train the models, ensuring robust performance evaluation and model parameter-tuning. For all model training, we used the Adam optimizer with a learning rate of 10^−4^, and the default hidden state dimension was set to 1024. Models were trained for 300 epochs. To fine-tune some models, we used the fine-tuning dataset to conduct further 50-epoch model training. For the ABCD data, we have *n*=95 subjects for fine-tune, and the fine-tuned models were evaluated in the independent test dataset (*n* = 285). For the HCP data, we have *n* =500 subjects in the fine-tuning set and *n* =565 subjects serving as an independent test. Model performance was assessed using standard metrics: Accuracy, Precision, and F1-score, calculated on the held-out test set. The results are presented in Table 3 & 4 (as referenced in the Results section).

## Code availability

Data and code used in this study are available at https://github.com/sbci-brain/BridgeBP.git, where applicable.

## Author contributions statement

Ziqian Zhang: Formal analysis; Investigation; Software; Validation; Visualization; Writing – original draft; Writing – review & editing. Alex Haopeng Liu: Writing – original draft; Writing – review & editing. Zhengwu Zhang: Conceptualization; Supervision; Writing – original draft; Writing – review & editing.

## Competing interests

The authors declare no competing interests.

## Notes

### Competing Interest Statement

The authors have declared no competing interest.

### Funding

- National Institute on Drug Abuse: 10.13039/100000026

