## Supplementary Material for "BridgeBP: A Toolbox for Bridging Brain Parcellations and Standardizing Structural Connectivity Matrices"

The following figures provide additional quantitative assessments of the BridgeBP conversion process, complementing the results presented in the main paper. To obtain a more comprehensive evaluation of reconstruction performance, we include (i) error-based metrics using mean squared error (MSE) as an alternative to correlation, (ii) scatter comparisons of true versus reconstructed non-zero structural connectivity (SC) across multiple atlases, and (iii) group-level analyses examining the effects of kernel smoothing on reconstruction performance. Figure S1 shows the standard deviations of conversion accuracy in Figure 3 (main text) for randomly selected 100 ABCD subjects.

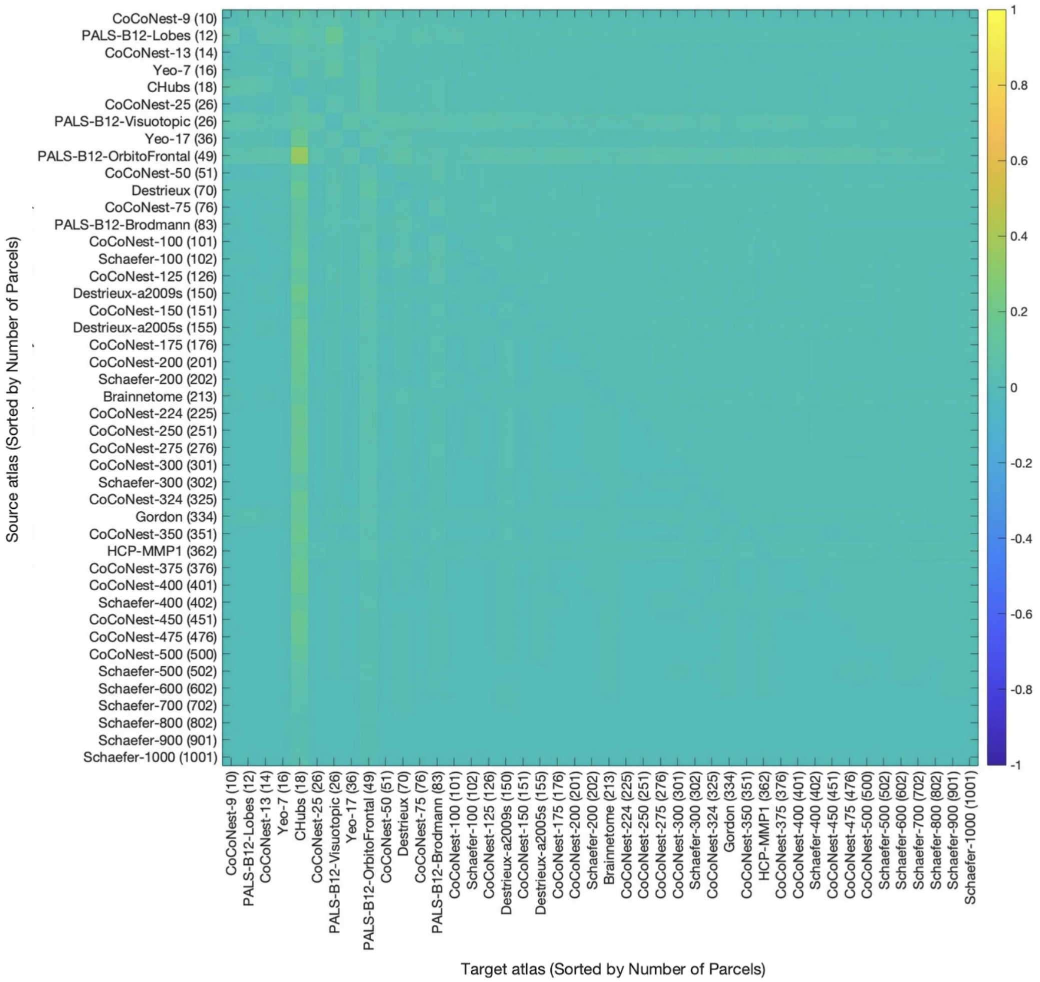


**Figure S1.** Standard deviations of conversion accuracy in Figure 3 (main text), reflecting small variability in SC conversion between atlas pairs.

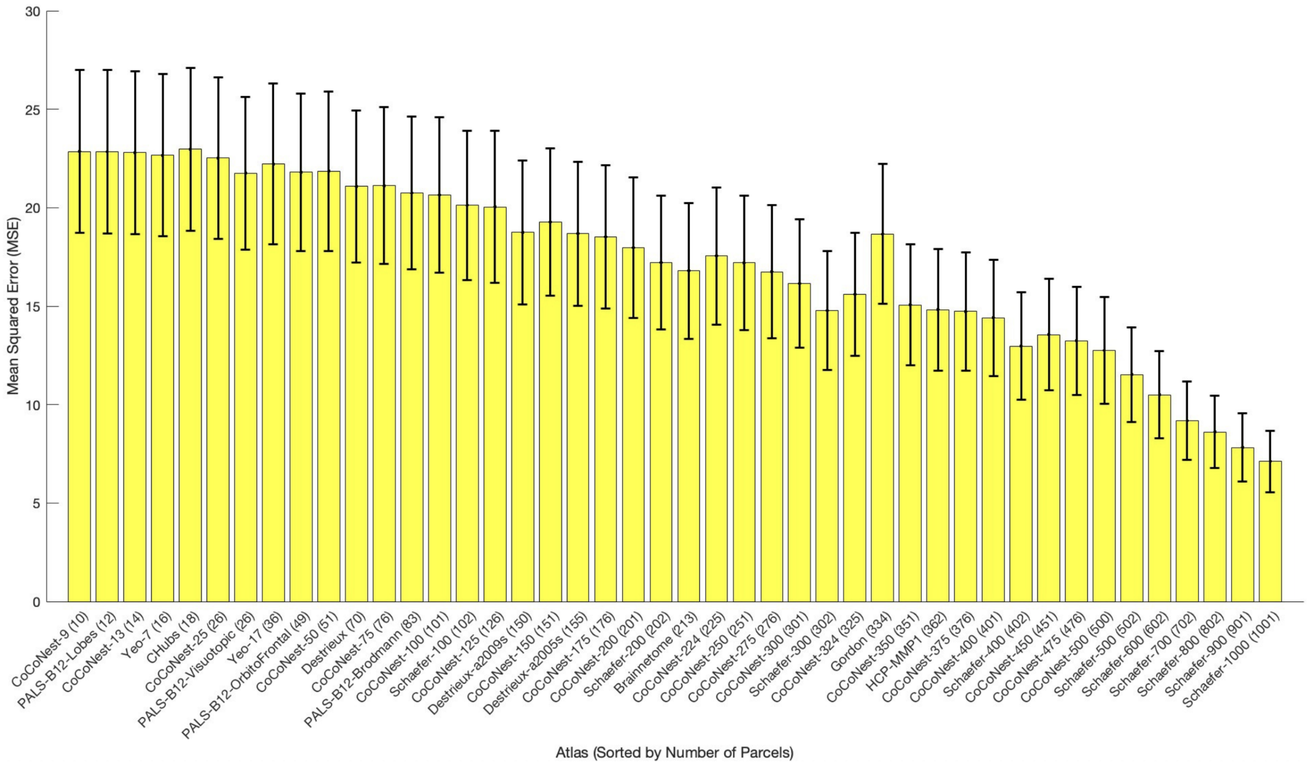


**(a)**

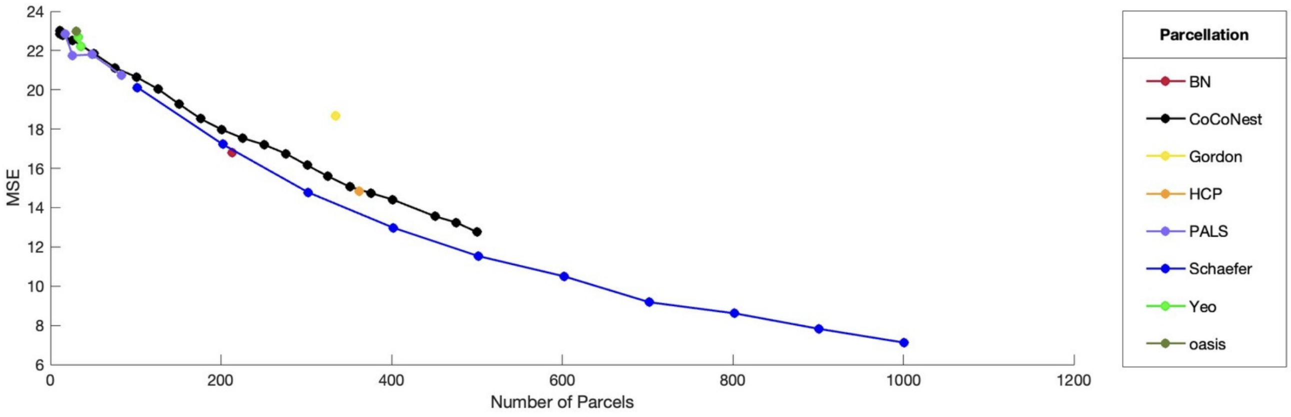


**(b)**

**Figure S2.** Similar to Figure 2 (main text), but measured using MSE. The Standard deviations in Panel (A) were computed based on 100 subjects.

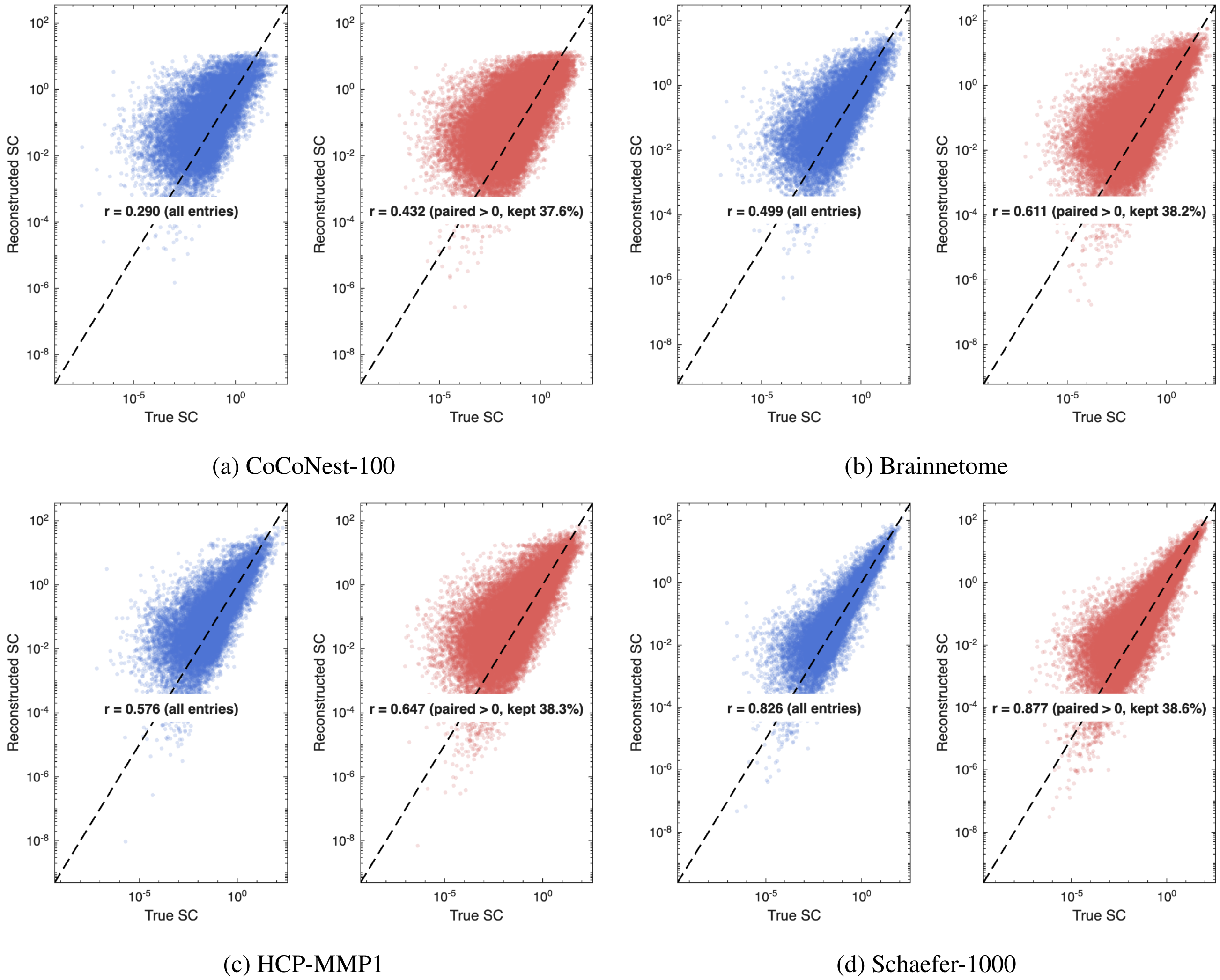


**Figure S3.** Scatter comparison of true versus reconstructed SC after zero-value removal. Results are shown for four atlas: (a) CoCoNest-100 (101 parcels); (b) Brainnetome (213 parcels); (c) HCP-MMP1 (362 parcels); (d) Schaefer-1000 (1001 parcels). Each panel illustrates the reconstructed SC values and the corresponding scatter plots between true and reconstructed matrices, with Pearson correlations ($r$) reported for all non-zero entries.

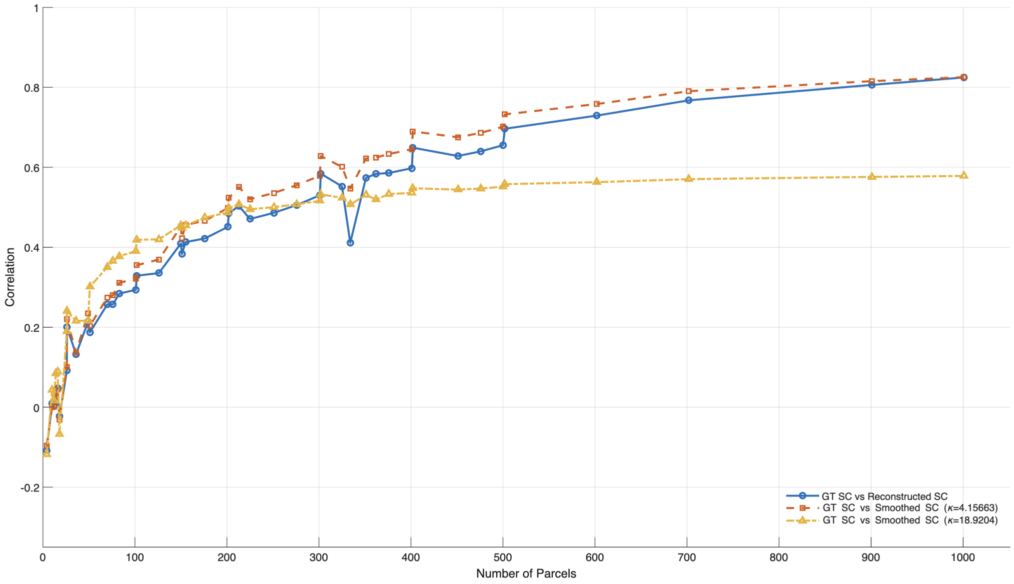


**Figure S4.** Group mean correlation between ground truth (GT) and reconstructed SC across 100 subjects. Correlations are shown as a function of the number of parcels, comparing GT SC with reconstructed SC (blue) and with smoothed SCs obtained using two different kernel bandwidths ($\kappa = 4.1563$ and $\kappa = 18.9204$).
